# Multinomial modelling of TB/HIV co-infection yields a robust predictive signature and generates hypotheses about the HIV+TB+ disease state

**DOI:** 10.1101/497842

**Authors:** Fergal J Duffy, Ethan G. Thompson, Thomas J. Scriba, Daniel E Zak

**Affiliations:** Center for Infectious Disease Research (formerly Seattle Biomedical Research Institute), Seattle WA, USA; South African Tuberculosis Vaccine Initiative, Institute of Infectious Disease and Molecular Medicine & Department of Pathology, University of Cape Town, Cape Town, South Africa

**Keywords:** Tuberculosis, HIV, co-infection, machine learning, multinomial, interferon, microarray, blood transcription

## Abstract

**Background:** Current diagnostics are inadequate to meet the challenges presented by coinfection with *Mycobacterium tuberculosis* (Mtb) and HIV, the leading cause of death for HIV-infected individuals. Improved characterisation of Mtb/HIV coinfection as a distinct disease state may lead to better identification and treatment of affected individuals.

**Methods:** Four previously published TB and HIV co-infection related datasets were used to train and validate multinomial machine learning classifiers that simultaneously predict TB and HIV status. Classifier predictive performance was measured using leave-one-out cross validation on the training set and blind predictive performance on multiple test sets using area under the ROC curve (AUC) as the performance metric. Linear modelling of signature gene expression was applied to systematically classify genes as TB-only, HIV-only or combined TB/HIV.

**Results:** The optimal signature discovered was a single 10-gene random forest multinomial signature that robustly discriminates active tuberculosis (TB) from other non-TB disease states with improved performance compared with previously published signatures (AUC: 0. 87), and specifically discriminates active TB/HIV co-infection from all other conditions (AUC: 0.88). Signature genes exhibited a variety of transcriptional patterns including both TB-only and HIV-only response genes and genes with expression patterns driven by interactions between HIV and TB infection states, including the CD8+ T-cell receptor LAG3 and the apoptosis-related gene CERKL.

**Conclusions:** By explicitly including distinct disease states within the machine learning analysis framework, we developed a compact and highly diagnostic signature that simultaneously discriminates multiple disease states associated with Mtb/HIV co-infection. Examination of the expression patterns of signature genes suggests mechanisms underlying the unique inflammatory conditions associated with active TB in the presence of HIV. In particular, we observed that disregulation of CD8+ effector T-cell and NK-cell associated genes may be an important feature of Mtb/HIV co-infection.

## Background

Almost ¼ of the global population is infected with *Mycobacterium tuberculosis* (Mtb) [1] and over 1,600,000 people succumbed to active tuberculosis disease (TB) in 2016 alone[2]. TB ordinarily requires at least six months of antibiotic treatment in order to remove all traces of the infection, with drug resistant strains requiring two years of intensive treatment[3]. Individuals with HIV/AIDS are at particularly high risk of active TB, up to 30 times higher than for HIV-individuals prior to the start of antiretroviral therapy(ART)[4]. This relative risk declines after the initiation of ART, but still remains 2-3 times higher than the general population, and the biological mechanisms underlying this increased risk remain unclear.

The current standard for diagnosis of active TB is microscopic or culture-based detection of *M. tuberculosis* bacteria in a patient-derived sputum sample. Sputum-based tests suffer from several major limitations, including the amount of time it takes to culture slow-growing TB, and the necessity of having sufficient TB bacteria in the sputum for detection. This is a further issue for children and TB cases in HIV+ patients[5] where low numbers of TB bacilli in the sputum may give a false-negative result. The Xpert MTB/RIF test[6] has enabled rapid TB diagnosis by detecting the presence of *M.tb*-specific DNA in sputum, but the sensitivity of this test is diminished in sputum culture-negative TB[7]. Sputum also represents a dangerous vector of infection for health-care workers analysing and handling sputum samples, due to the potential presence of live *M. tb* bacteria[8]. New TB diagnostic methods that do not necessitate the detection of large numbers of TB bacilli in sputum are therefore critically required to serve populations at high risk of TB. Blood-based signatures are an attractive alternative, as blood is a clinically accessible readout of the immunological state of the body.

Whole blood gene expression signatures that are diagnostic for TB have been described in many previous studies[9–12], but these signatures are generally focused on a single binary comparison, e.g. latent TB vs active TB or active TB vs other diseases. In this study, we develop multi-class multinomial signatures that explicitly model the TB and HIV state of each patient. Our analysis integrates published data from several cohorts and evaluate a range of machine-learning approaches to generate a multinomial model that specifically discriminates TB from non-TB disease states while simultaneously discriminating HIV+ TB as a unique disease state.

## Methods

All computational and statistical analyses were performed using the R language for statistical computing[13].

### Microarray Normalisation, Probe Filtering and Data Preparation

Microarray data from four TB/HIV cohorts were downloaded from GEO: GSE37250, GSE39941, GSE19491, GSE42834. The precise sample compositions of each of these datasets are provided in Table 1 (GSE37250) and Table 3 (GSE39941, GSE19491, GSE42834).

**Table 1:** Adult Training and Test Set

|  | Active TB |  | Latent TB |  | Other Diseases |  | TOTAL |
| --- | --- | --- | --- | --- | --- | --- | --- |
|  | HIV+ | HIV- | HIV+ | HIV- | HIV+ | HIV- |  |
| Malawi | 51 | 51 | 36 | 35 | 30 | 34 | 237 |
| South Africa | 47 | 46 | 48 | 48 | 62 | 49 | 300 |

Microarray datasets were downloaded in the form of GEO series matrix files, background subtracted and quantile normalized. All reference and variable probe selection was performed using the GSE37250 Malawi adult data. 20 reference probes were selected by eliminating any probe with any expression value below the absolute value of the smallest expression value observed in the dataset. References were then selected as the 20 remaining probes with the smallest inter-quartile range (IQR) of expression in the dataset.

To pre-select probes likely to be discriminatory for TB, only probes with an IQR of above 1.5 (log2 normalised expression) were kept. This resulted in 554 candidate model-variable probes to be used in the model. Additional File 1 lists the variable and reference probes used in this work.

Before model training, each sample in every dataset was normalised by calculating the mean expression level for the reference probes, and subtracting this mean reference level from each model-variable probe. The GSE19491 dataset, which was measured using the Illumina 12v3 platform as opposed to the Illumina 12v4 platform used for the other datasets, was missing 4 of the 20 selected reference probes. The remaining 16 probes alone were therefore used to normalise this dataset.

### Machine-Learning Model Training, Feature Shrinking and Model Selection

Five distinct machine-learning algorithms were used to train predictive models on the adult South African dataset, using the R caret[14]package. These were:

1. Random Forest (RF) (R randomForest package[15]), is an algorithm based on training an ensemble of decision trees using randomly split subsets of the training samples and training variables, all of which then ‘vote’ to classify new samples.
2. Support Vector Machine (SVM) using RBF kernel (R kernlab[16] package). SVMs attempt to find the optimal linear hyperplane decision boundary separating the two classes in n-dimensional space, where n is the number of features the SVM is trained on. The RBF, or Radial Basis Function kernel projects this n-dimensional feature space into a higher dimension to allow the identification of a linear decision boundary in a higher dimensional space if one cannot be found in the input n-dimensional space.
3. Neural Networks (NN) (R nnet[17] package). NNs are comprised of a network of input nodes (1 per-feature), connected to output nodes (1 per possible outcome) via one or more ‘hidden’ layers of nodes. Each node represents a logistic regression function, based on the input value, and the weight given to each input node (which in turn determines the output classification) is determined during training.
4. Elastic-net Logistic Regression (R glmnet[18] package), is a form of logistic regression with regularisation of the linear coefficients applied to control overfitting.
5. K-Nearest Neighbor (KNN) (R caret[14] package), classifies samples by determining the ‘k’ most similar samples by Euclidian distance between sample genes and having them ‘vote’ on the classification.

All of these algorithms can be trained to produce binary (exactly 2 distinct classes, such as TB vs LTB) or multinomial (more than 2 classes) classifier models.

In this study, each algorithm was trained using normalised microarray data comprising all 554 most-variable probes (selected as described above) on each of four subsets of samples of the adult training data. These subsets were 1) the entire dataset, including TB, latent TB (LTB) and other disease (OD) samples including HIV+ and HIV- samples, 2) All TB and LTB samples including HIV+ and HIV-samples (i.e. OD excluded), 3) All HIV+ TB and LTB samples, (i.e. all HIV- and OD excluded), 4) All HIV-TB and LTB samples (i.e. all HIV+ and OD excluded). Models were trained to simultaneously predict both the TB and HIV status of each training sample, i.e. models trained on subset 1) explicitly classified samples as one of 6 classes: TB:HIV+, TB:HIV-, LTB:HIV+, LTB:HIV-, OD:HIV+, or OD:HIV-; models trained on subset 2) classified samples as one of 4 classes TB:HIV+, TB:HIV-, LTB:HIV+, or LTB:HIV-, and models trained on subsets 3) and 4) were binary models that classified samples as TB or LTB only, as HIV status was constant in these subsets.

After models were trained on the initial 554 probes, the models were sequentially shrunk to obtain probe-reduced models that only comprised the most important 250, 50, 25, 15, or 10 probes from the initial set. Probe importance rankings were calculated using the *varImp* function supplied by the caret package. This function implements algorithm-specific methods for evaluating how much each probe contributes to the classification performance of the model. In the case of random forests, the difference in out-of-bag error[19] with and without the inclusion of a single probe was used to rank the probes in order of importance. For Elastic-net logistic regression models, the probe variable coefficient was used to rank the probes. For Neural Networks, Garson’s algorithm[20] was used to calculate probe importance from network weights. For Random Forests, probe importance was measured as the difference in predictive performance comparing all trees that contain the probe with trees that lack that probe. For the remaining modelling approaches (SVM, KNN), the univariate predictive power of the individual probe was used to rank importance in an algorithm-independent way.

For each algorithm-subset combination, the most important 250 out of the original 554 probes were selected, the model re-trained on this subset of probes, and this probe-reduced model used to predict the left-out sample. This procedure was then repeated to iteratively shrink each model to contain the 50, 25, 15 and 10 most important probes from the previous step. This entire process was carried out for each held-out sample in the cross validation so that the sequential shrinking and prediction steps were performed independently for every held out sample.

Training performance was assessed using leave-one-out cross validation (LOOCV). Initially, a single sample from the overall training set was held out, then a classifier model was trained on the remaining samples, and used to predict the status of the held-out sample. This procedure was repeated for every training sample, and model performance was then calculated using predictions on the held-out samples. Areas under the ROC curves (AUCs) for model discrimination of TB vs non-TB samples in the training subset were calculated for predictions on the held out samples, and this was used as the performance metric for initial model structure selection. For models that predicted more than 2 classes, TB predictions were calculated as the sum of TB-related prediction classes, e.g for 6 class models the overall TB prediction value was calculated as the TB:HIV+ prediction value plus the TB:HIV-prediction value.

Only “small” models (i.e., those consisting of 10, 15 or 25 probes) were considered for application to the test sets. To choose the algorithm/class-complexity/training set between these 3 probe sizes, initially the 10-probe model was selected. If either the 15 or 25-probe model showed significantly better LOOCV performance on the training set, that model was used. Significance was evaluated by comparing ROC AUCs using the *roc.test* function from the pROC[21] R package with a threshold of p<0.05.

### Model Predictions on New Datasets

After models were selected by recursive LOOCV evaluation as described above, each selected model was retrained using the most-commonly selected features from the LOOCV and re-parameterised on the entire relevant training data subset. Predictions were made using the *predict* function from the caret package to calculate class probabilities, and prediction accuracies were assessed by calculating TB vs non-TB ROC curves using the R pROC package. As described for the LOOCV procedure above, in the case ofmodels that predicted more than 2 classes, TB predictions were calculated as the sum of TB-related prediction classes, e.g for 6 class models the overall TB prediction value was calculated as the TB:HIV+ prediction value plus the TB:HIV-prediction value. Performance of the new signatures on the test sets was benchmarked against the performance of two previously-reported TB gene signatures: the three gene multi-cohort diagnostic signature developed by Sweeney et al[27], which we term the ‘threeGene’ signature; and our 16-gene correlate of TB risk [23], which we term the ‘ACS’ signature (referring to the Adolescent Cohort Study from which the signature was derived). For predictions using the threeGene signature, datasets were downloaded in raw non-normalised format from GEO before being quantile normalised and baseline corrected using the log-exponential method using the R limma package[22]. The threeGene score was then directly calculated as (GBP5 + DUSP3)/2 – KLF2. For the ACS model predictions, datasets were prepared and normalised and scored as described in[23].

### Linear modelling of disease state

Linear regression models were fit to signature genes in order to assess the contribution of TB and HIV status to gene expression. Expression of each gene was fit to a linear regression model (R *lm* function) of the form:

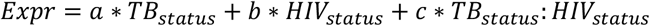

TB and HIV status were encoded as binary variables with 1 meaning active TB/HIV+ and 0 meaning latent TB/HIV-. Non-TB other disease samples were excluded from this analysis. The p-value of the model coefficients a, b and c were calculated using the R *summary.lm* function, and a false discovery rate correction applied.

### Gene set enrichment analysis

Gene set enrichment analysis was performed using the R tmod[24] package, using the blood transcriptional gene sets previously described by Li et al[25] and Chaussabel et al[26]. P-values were calculated using the hypergeometric test as implemented in the tmodHGtest function, using all included microarray gene symbols as the background.

## Results

### Development, cross-validation and selection of multinomial machine learning models for predicting TB and HIV

We used data from a previously-published cohort [9] of 537 adults from Malawi and South Africa, comprised of samples from individuals diagnosed with active tuberculosis (TB), latent tuberculosis (LTB) or other non-TB diseases with clinical symptoms consistent with TB (OD). Roughly half of these individuals were also HIV+ (Table 1). These transcriptional profiles were used to develop and test multinomial machine learning approaches to specifically identify each symptomatic subset.

Machine learning models were trained on the South African adult dataset described in Table 1, with the Malawian adults used as an independent test set. In order to focus on the strongest signal probes, an initial down-selection step was performed where only probes with a log2 normalised expression interquartile range of at least 1.5 in the South African set were considered for model training (554 probes).

Models were trained to classify all or relevant subsets of the data into 2 (binary classifier), 4 (multinomial), or 6 (multinomial) classes, using a diverse panel of machine-learning algorithms. Two-class models were trained to classify a sample as either active or latent TB. Two different two class models were trained for each algorithm, one on HIV-TB and LTB samples only, and another on HIV+ TB and LTB samples only. Four-class models were trained to classify a sample as active or latent TB and as HIV+ or HIV-simultaneously, using all TB and LTB samples, both HIV+ and HIV-. Six-class models were trained to classify a sample as active TB, latent TB or other disease, and as HIV+ or HIV-, and were trained on the entire dataset, including TB, LTB and OD, both HIV+ and HIV-. Machine learning algorithms used were Random Forests, Neural Networks, Support Vector Machines, Elastic-Net Logistic Regression, and k-Nearest Neighbours.

Initially, each model was trained using all 554 pre-selected probes. Starting from this initial model, the most important model probes were selected and the models recursively shrunk to use smaller numbers of probes (see Methods). Leave-one-out cross validation (LOOCV) performance on the training set was evaluated by measuring area under the ROC curve (AUC). Figure 2 shows the results of the LOOCV and recursive shrinking for each algorithm (Figure 1). LOOCV AUCs are uniformly strong, almost all above 0.8. LOOCV performance of multinomial 4- and 6- class models is similar to that of the binary classification models.

**Figure 1:**
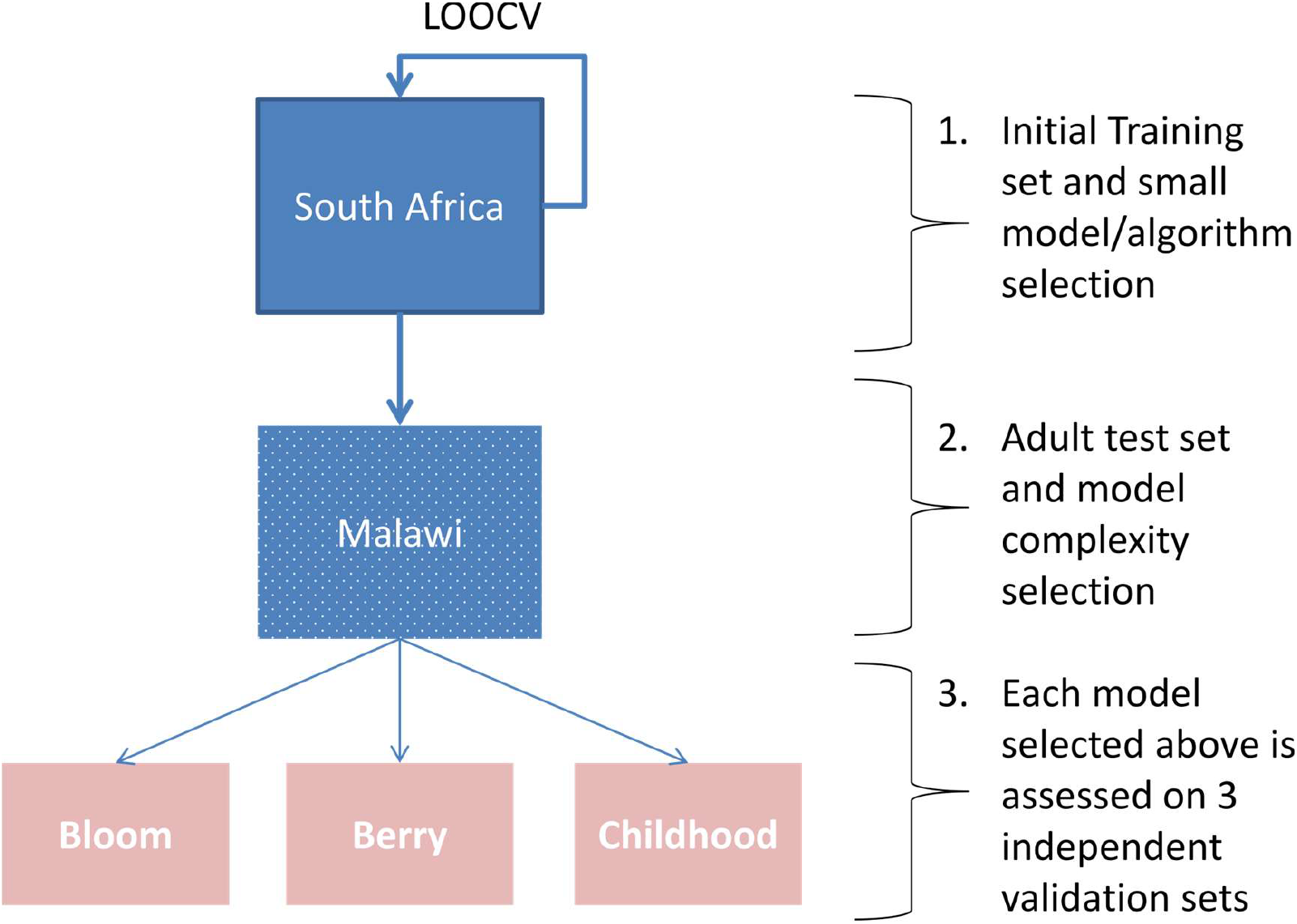
Analytical Plan. Outlines the model training, selection and prediction steps for the overall analysis

**Figure 2:**
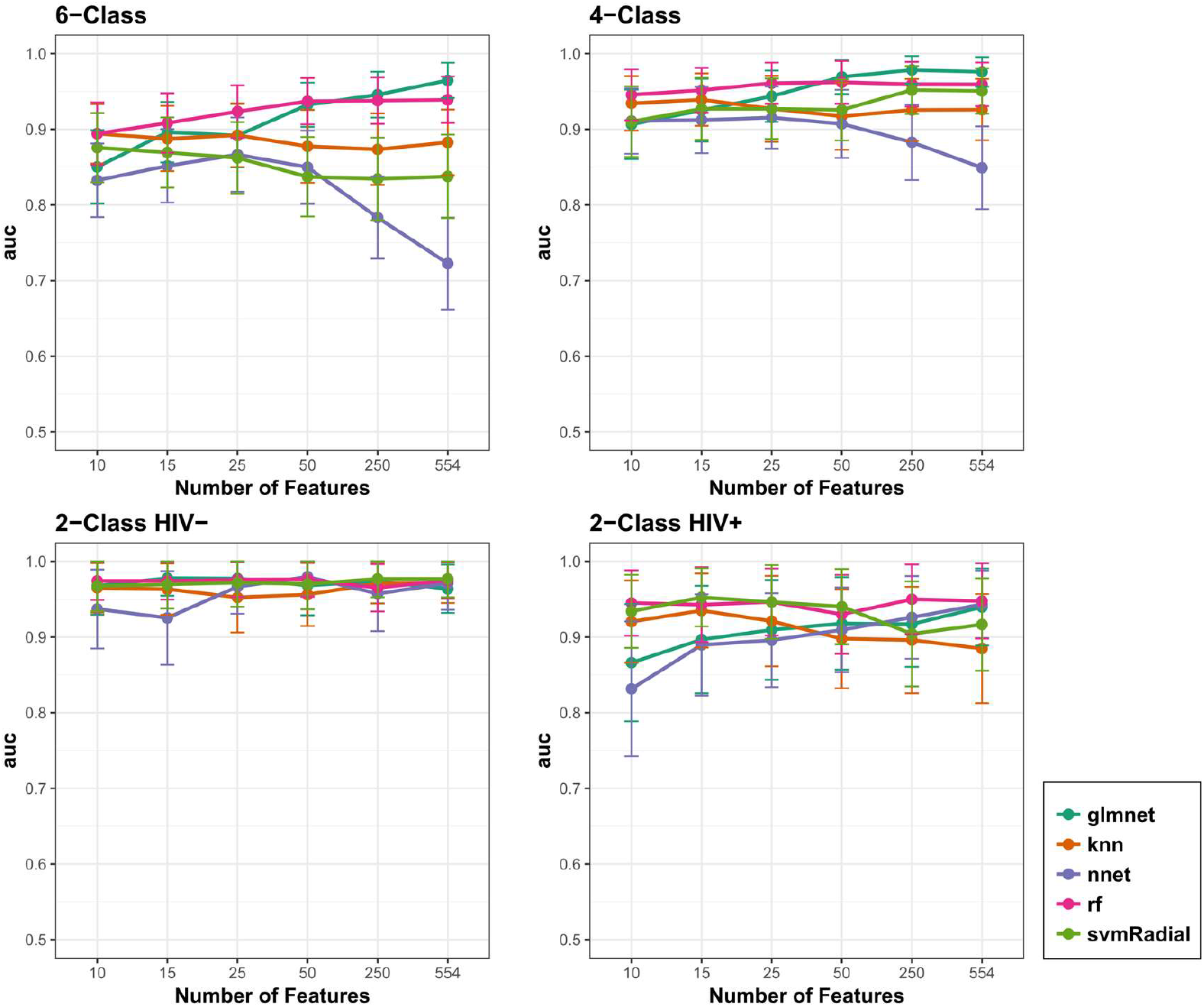
Training Cross Validation Results on Adult TB Samples. Leave-one-out cross validation results for models trained on South Africa adult data. Each panel plots the area under the ROC curve for five machine learning algorithms (glmnet: Elastic-Net logistic regression, knn: k-Nearest Neighbours, nnet: Neural Network, rf: Random Forest, svmRadial, Support Vactor Machine with Radial Basis Function kernel) starting with models trained using all 554 probes, and iteratively shrunk to models trained on 10 probes only. Models were trained to classify the data into 6 (TB:HIV+, TB:HIV-, LTB:HIV+, LTB:HIV-, OD:HIV+, OD:HIV-), 4 (TB:HIV+, TB:HIV-, LTB:HIV+, LTB:HIV-) and 2 (TB, LTB) classes. Two types of 2-class models were trained: using all HIV+ or all HIV-samples. Error bars show bootstrap-estimated 95% confidence intervals around the AUC.

An ideal model shows high predictive performance based on a small number of interpretable genes. A set of small models for further analysis were chosen by initially selecting the smallest (10 probe) model for each algorithm and classification complexity, and only selecting a larger model if it showed significantly stronger LOOCV performance. As performance, illustrated in

Figure 1 (a) and (b), was largely uniform across model sizes, the 10 probe models were universally selected. Table 2 lists the training cross-validation performance of each of these models, in terms of their area under the ROC curve. From Table 2, in three of the four complexity cases for the South Africa training set, Random Forest was the highest-performing algorithm, thus Random Forest models were selected for all further analyses.

**Table 2:**
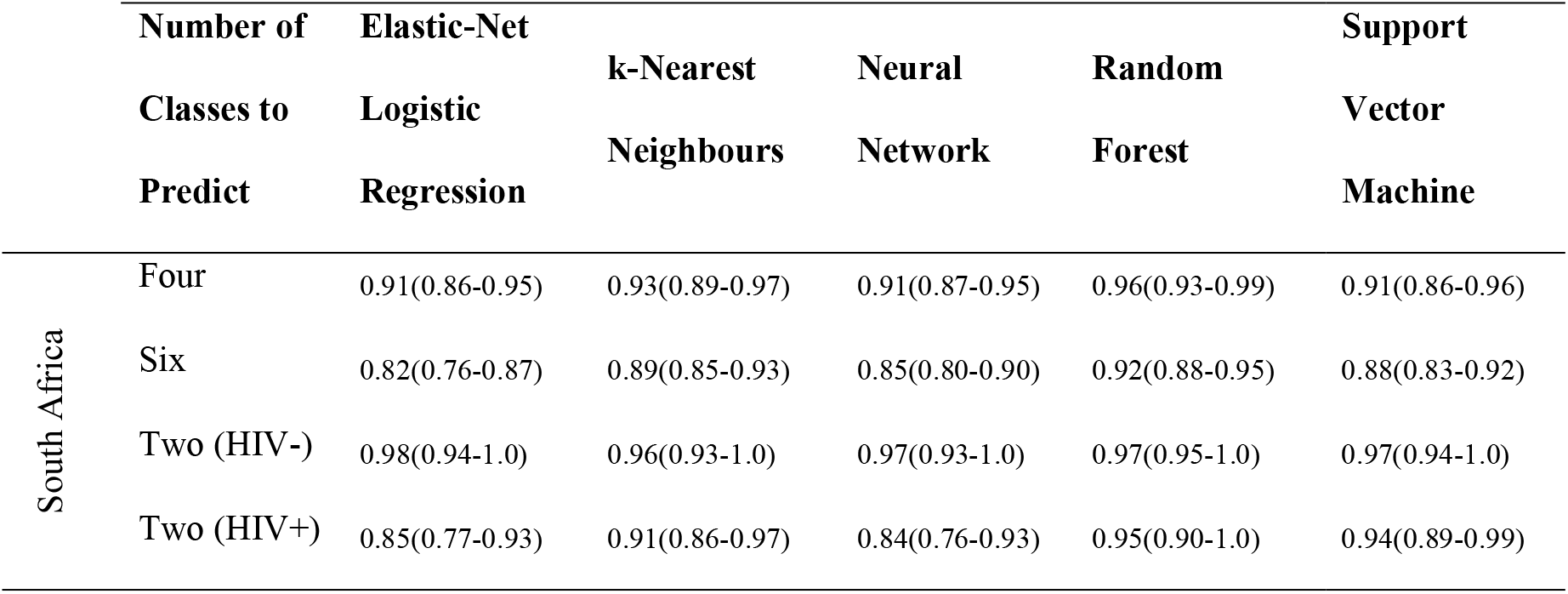
Top LOOCV Model Structuresy.

|  | Number of<br>Classes to<br>Predict | Elastic-Net<br>Logistic<br>Regression | k-Nearest<br>Neighbours | Neural<br>Network | Random<br>Forest | Support<br>Vector<br>Machine |
| --- | --- | --- | --- | --- | --- | --- |
| South Africa | Four | 0.91(0.86-0.95) | 0.93(0.89-0.97) | 0.91(0.87-0.95) | 0.96(0.93-0.99) | 0.91(0.86-0.96) |
|  | Six | 0.82(0.76-0.87) | 0.89(0.85-0.93) | 0.85(0.80-0.90) | 0.92(0.88-0.95) | 0.88(0.83-0.92) |
|  | Two (HIV-) | 0.98(0.94-1.0) | 0.96(0.93-1.0) | 0.97(0.93-1.0) | 0.97(0.95-1.0) | 0.97(0.94-1.0) |
|  | Two (HIV+) | 0.85(0.77-0.93) | 0.91(0.86-0.97) | 0.84(0.76-0.93) | 0.95(0.90-1.0) | 0.94(0.89-0.99) |

### A six-class multinomial model outperforms previously published signatures for identifying active TB in several independent test sets

The Malawian adults were used as an independent test set for the selected models.

Figure 3 (A) shows ROC curves representing the predictive ability of South Africa-derived models to specifically identity HIV- and HIV+ active TB samples vs latent TB and other diseases. These models were accompanied by two previously-reportedTB signatures: our 16-gene correlate of TB risk [23], termed here as the ‘ACS’ signature, and the three-gene multi cohort diagnostic signature developed by Sweeney et al[27], termed here as the ‘threeGene’ signature. The multinomial six-class Random Forest model outperformed all other models (AUC: 0.88, sensitivity 80%, specificity 82%), although performance of the threeGene model was very similar (AUC: 0.87 vs AUC 0.88).

**Figure 3:**
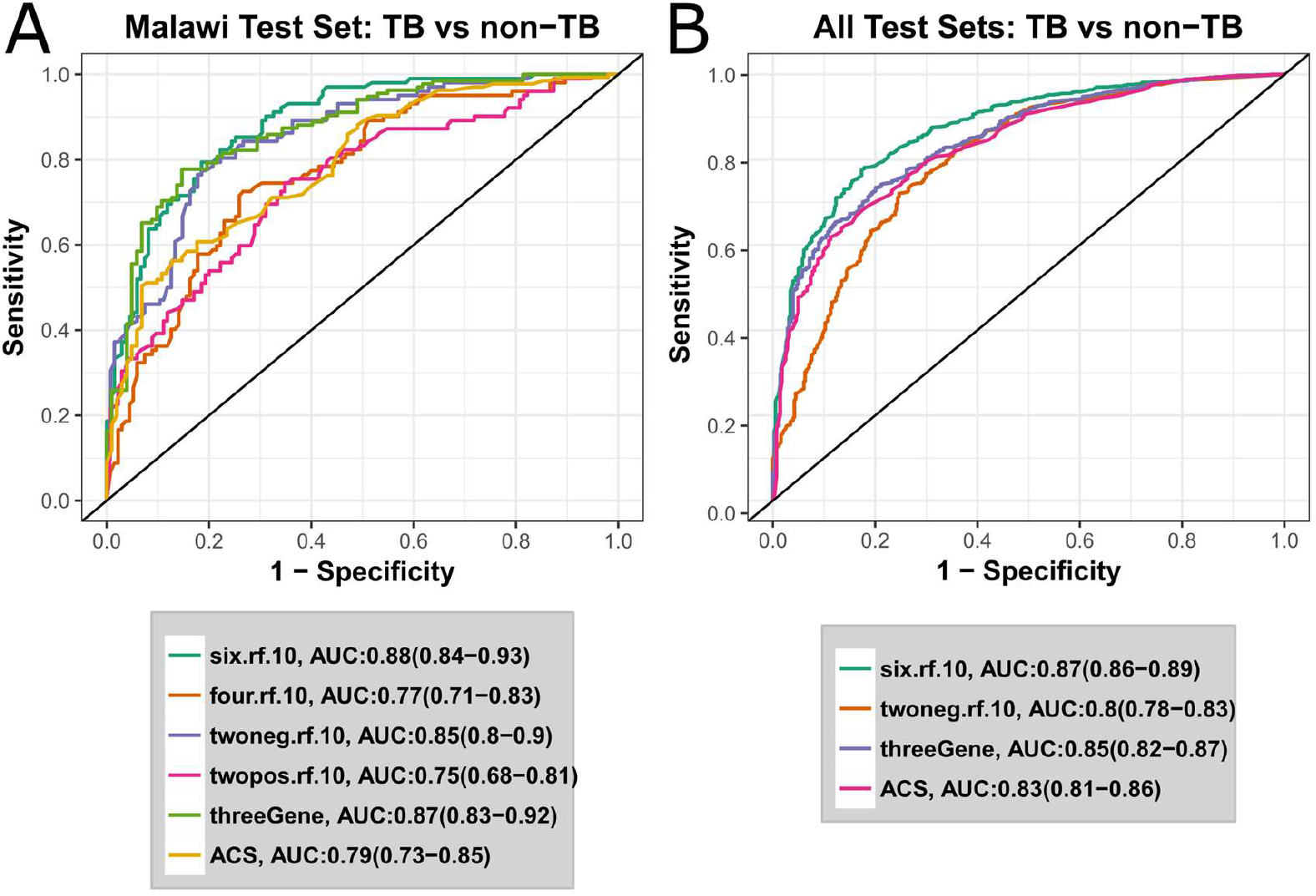
A six-class multinomial model optimally predicts 4 independent test sets. ROC curves for active TB vs non-TB classification of independent test sets. Legends shows the AUC for each model, with the 95% confidence intervals in parentheses. Models developed in this study are named in the form <number-of-classes>.<algorithm>.<number-of-probes>. E.g. six.rf.10 is the 10 probe random forest model trained to predict 6 classes. twoneg and twopos refer to 2-class models trained on HIV- or HIV+ samples respectively. threeGene refers to the signature described by Khatri et al[32], and ACS refers to the signature described by Zak et al[23]. **A** ROC curves for classification of the Malawi test samples from the Kaforou cohort. **B** ROC curves for Malawi test set plus the three further independent test sets described in Table 3

**Table 3:**
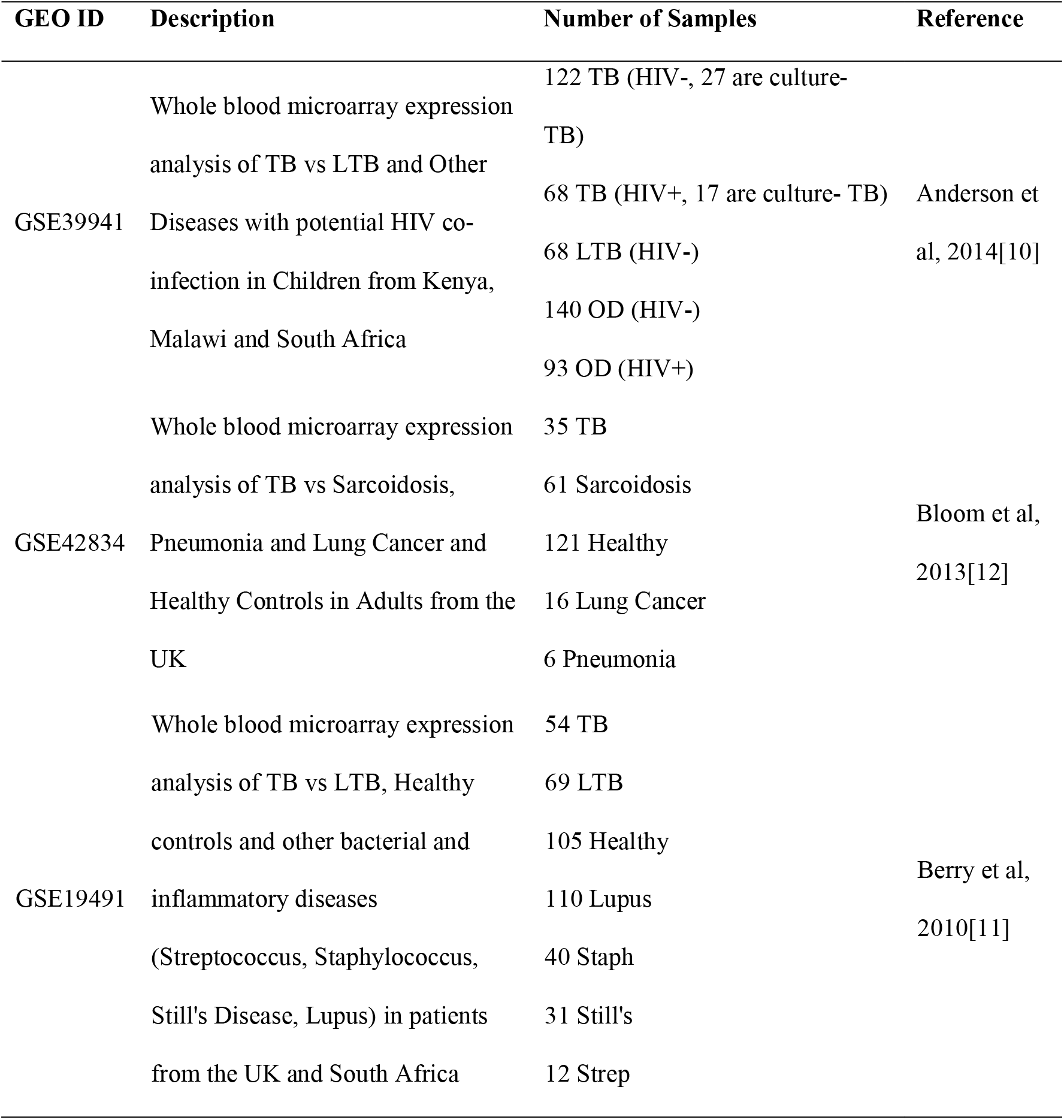
Further Validation Datasets

To more thoroughly validate the six-class multinomial model, classification performance was evaluated using three additional previously-published whole-blood microarray datasets[10–12]. To reduce technical sources of variability as much as possible, test sets were selected that used the same Illumina HumanHT-12 microarray platform as was used for the Kaforou cohort. These additional test sets comprised a broad range of samples, including adult and childhood TB; TB vs other inflammatory, bacterial and pulmonary diseases; and samples taken from a range of geographical locations (Table 3). The two-class HIV-model was also included as a comparator for the six-class multinomial model.

Figure 3 (B) shows the ROC curves for the 10-gene six-class model, the 10-gene binary model and the two previously-described external signatures. Again, the six-class 10-gene signature was the overall top performer (AUC 0.88, sensitivity 80%, specificity 82%). This is significantly better predictive performance than the top external model, the threeGene signature (p=0.006 by a single tailed DeLong[28] test). Thus, multinomial modelling of TB disease states significantly improved the accuracy of discrimation of TB vs. non-TB samples.

### The 10-gene multinomial signature identifies HIV+TB as a distinct disease state

To further evaluate the performance of the 10-gene six-class signature, particularly the potential of this signature to perform multi-class discrimination, we tested whether it can specifically identify HIV+ TB samples from all other samples in the Malawi test set (Figure 4 (A)). The signature accurately discriminated HIV+ TB from HIV-TB samples (AUC: 0.88) and HIV+ TB samples from all other samples (HIV+ TB, HIV-/+ latent TB, HIV-/+ other diseases, AUC: 0.86). This result suggests that HIV+ TB may exist as a distinct transcriptional state. Box- and dot-plots of normalised expression for the 10 genes in the multinomial signature for active and latent TB individuals in the combined South-African and Malawian cohorts, stratified by TB and HIV status reveal a diverse pattern of transcriptional responses (Figure 4 (B)).

**Figure 4:**
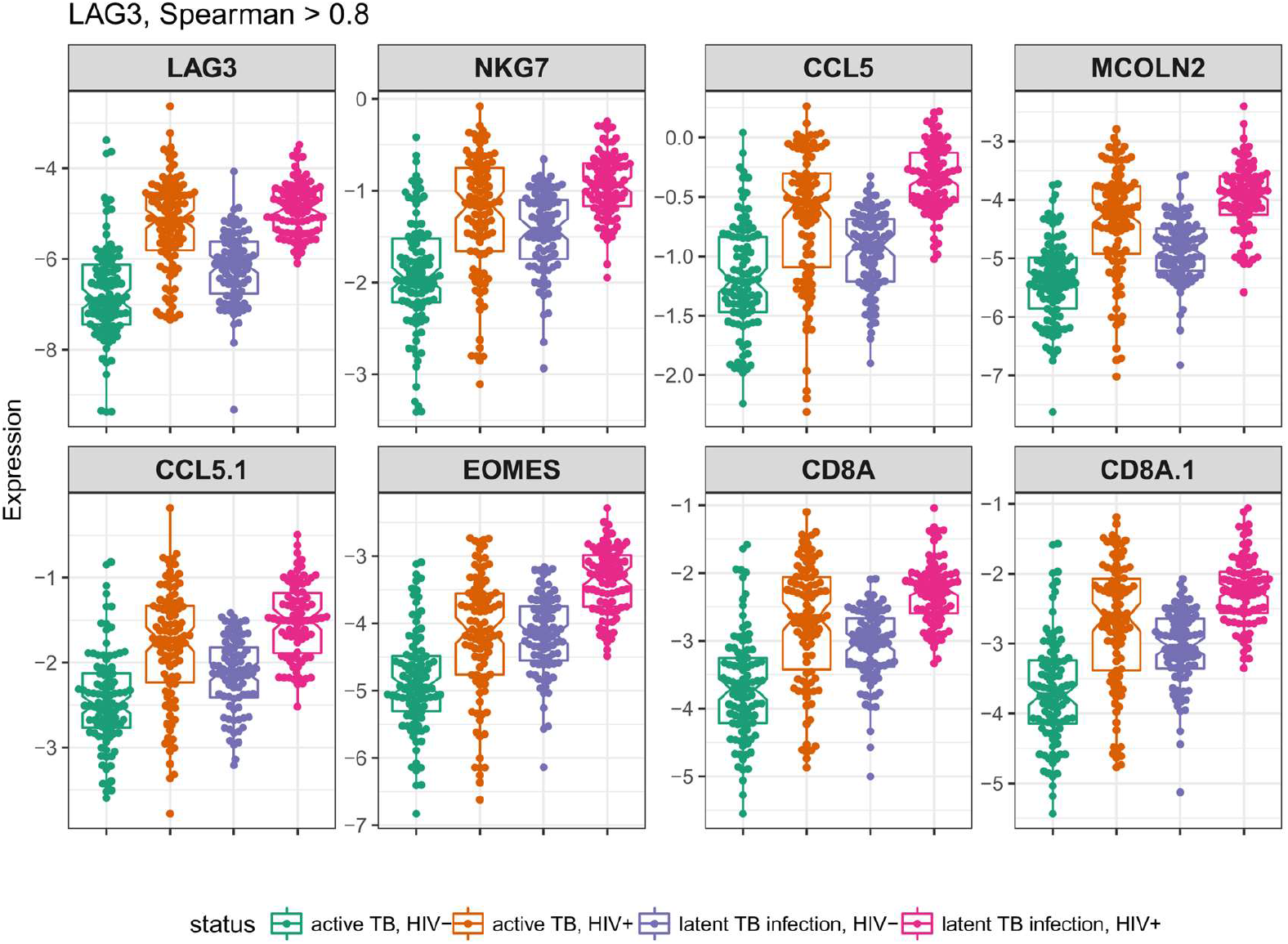
The six-class multinomial model identifies HIV+ TB as a distinct state. **A** ROC curves for the 10-gene six-class multinomial model discriminating HIV+ active TB samples from HIV-active TB samples, and HIV+ active TB samples from HIV-active TB and HIV+/- LTB and HIV+/- OD samples in the Malawi test set. **B** Dot- and boxplots of expression levels of six-class multinomial model genes in the entire Kaforou dataset. **C** Six-class multinomial genes classified by their TB/HIV behaviour as determined by fitting linear models to gene expression as a function of disease state. TB upregulated genes are indicated in orange and downregulated genes shown in blue

Using HIV-latent TB samples as a baseline, 8 of the 10 signature genes are either downregulated in both active TB and HIV+ patients, or upregulated in both active TB and HIV+ patients. Two exceptions are LAG3 and CERKL. To more quantitatively determine the transcriptional patterns of these genes, linear models with gene expression as a function of TB and HIV status were fit, including a TB:HIV interaction term. Genes with significant (FDR<0.01) TB or HIV model coefficients were identified as TB- or HIV-independent disease signature genes, and genes with both TB and HIV significant coefficients were identified as overlapping signature genes (Figure 4 (C)). Three of the ten genes in the signature were independent disease signature genes, with CD160 being the sole HIV-specific gene and CD36 and ZDHHC19 as the only TB-specific genes. The largest group of signature genes exhibits a unidirectional additive expression pattern, either downregulated in both TB and HIV (CD40LG, ID3) or upregulated in both TB and HIV (GBP6, C1QB). Interestingly, the CD8+ immune checkpoint gene LAG3 is upregulated in HIV+ individuals but downregulated in active TB. Two genes exhibit a significant interaction term: FCGR1B and CERKL. This interaction suggests crosstalk between the TB and HIV transcriptional response. In the case of FCGR1B, transcription reaches a saturated level in HIV-active TB that is not exceeded in HIV+ active TB. As FCGR1B is a cell surface receptor specific to macrophages, monocytes and neutrophils [EBI Expression Atlas, www.ebi.ac.uk/gxa], this saturation point may correspond with a maximum surface density of receptor or a maximal blood concentration for these cell types. CERKL, a negative regulator of apoptosis caused by oxidative stress[29], shows a more complex regulatory pattern where HIV-active TB is upregulated compared to all other states.

### Biological pathways associated with divergent TB/HIV expression patterns reveal HIV+ TB as a distinct disease state

Analysis of the genes comprising the 10-gene six-class signature identified LAG3 and CERKL as exhibiting distinct expression patterns. As the signature genes reflect a minimal set of genes necessary to classify disease states, we hypothesised that there may be other genes closely correlated with LAG3 and CERKL that could shed light on the biological processes driving the opposing regulation they exhibit.

The expression of LAG3 was correlated very tightly (Spearman ρ > 0.8, p < 1e-32), with a set of eight genes similarly downregulated in active TB and upregulated in HIV (Table S2, Figure 5). Mapping these genes to blood transcriptional genes sets[25,26] revealed significant enrichment for cytotoxic T-cell and NK-cell pathways (Table S3), suggesting that dysregulation of immune effector cells is a distinguishing characteristic of HIV+ active TB when compared to HIV-TB, or HIV+ latent TB.

**Figure 5:** LAG3-correlated genes. Dot and boxplots for each microarray primer, named as the corresponding gene, strongly correlated with LAG3 (spearman correlation ρ>0.8) for latent and active TB samples from the Kaforou dataset.

In contrast, CERKL does not show similarly strong correlations (Spearman ρ>0.8) with any individual gene. At a more permissive correlation threshold (ρ>0.6), CERKL is correlated with 9 genes (Table S4), but this set of genes does not show significant enrichment for any gene set. Genes most strongly correlated with CERKL are the ribosomal-RNA processing gene HEATR1 and the ubiquitin ligase TRIM13.

## Discussion

In a clinical setting, a major challenge faced regarding TB diagnosis is to discriminate active TB from other diseases presenting with similar symptoms. The ROC curves shown in Figure 3 shows that the 10 gene six-class signature identified in this work significantly improves on existing signatures for identifying active TB in a wide variety of contexts, i.e. active TB vs healthy samples, active TB vs latent TB and active TB vs other diseases, with or without the presence of HIV co-infection. A major advantage of the meta-analytical approach taken here is the testing of each signature on a combination of cohorts at once. While ROC analysis can reveal the optimal classification performance on a single cohort, it is still necessary to choose an operating point or threshold to transform a continuous score into a dichotomous classifier. It is possible for a predictive signature to show a high sensitivity and specificity on many individual cohorts separately, but fail to recreate this performance when samples from all cohorts are combined. This is due to the signature score potentially having a differing optimal classification threshold on each cohort and will not be revealed by separate ROC analysis of each cohort. By combining cohorts, signatures with a stable “global” operating score are revealed. Thus, it can be seen in Figure 3 (B) that the across every cohort, the ten-gene six-class signature predicts with a sensitivity of 80% and specificity of 78%.

Explicit modelling of each cohort disease group has allowed us to hone in on transcriptional processes that specifically distinguish HIV+ active TB from HIV-active TB. Characterisation of a specific transcriptional state for HIV+ TB would improve understanding of how HIV increases TB risk, as well as illuminating on essential elements of an effective host response to TB missing from HIV+ TB patients. This analysis identified the CD8+ inhibitory checkpoint receptor LAG3. Linear modelling reveals significant upregulation of LAG3 in HIV infection, but also significant downregulated of LAG3 in active TB compared with latent TB (Figure 4 (B)). Upregulation of LAG3 is known to suppress T-cell activity in chronic HIV infection[30], and these exhausted T-cells show impaired production of the cytokines such as IL-2, IFNγ, and TNF, associated with an effective host response to TB[31]. LAG3 expression is also closely correlated with genes including the CD8A receptor; the CD8+ T-cell secreted chemokine CCL5/RANTES; the NK-cell granule gene NKG7, the CD8+ differentiation transcription factor EOMES, and the lysosomal membrane protein MCOLN2. All of these genes are involved in CD8+ or NK-cell effector activities. Thus, further investigation of a key signature gene has revealed that both innate (NK cell) and adaptive (CD8+ T-cell) effector function appears to be suppressed in HIV+ active TB relative to HIV-active TB, suggesting at least one mechanism for increased TB risk in HIV+ individuals.

Interestingly, another CD8+ T-cell inhibitory checkpoint receptor, CD160, was also selected as a signature gene. CD160 shows a similar pattern of expression to LAG3: upregulated in HIV+ patients, but downregulated in active TB. However, this downregulation in active TB is much less pronounced than for LAG3, and the TB coefficient was not found to be significant in linear modelling (FDR=0.08).

Overexpression of CERKL has been shown to protect cells from apoptosis while under oxidative stress [29]. The expression pattern of CERKL, which shows lower expression in HIV+ active TB compared to both HIV-active TB and HIV+ latent TB indicates that HIV/Mtb co-infected patients may have impaired protection against cellular death due to oxidative stress. This expression pattern hints at a complex balance of apoptotic signalling in HIV/TB co-infection that does simply mirror the interferon-driven inflammatory response.

## Conclusions

We have identified a broadly applicable active TB-specific 10-gene multinomial signature by validating candidate signatures with successively harder problems: training a diverse panel of candidate models on an adult test set; making blind predictions an independent adult test set from a different geographical cohort, albeit from the same study; making blind predictions on the combination of the adult test set with three additional independent cohorts; and finally testing for discrimination of HIV+ TB from HIV-TB.

While the signature shown here does not reach the diagnostic sensitivity required to be a practical alternative to sputum culture for clinical use (>98% sensitivity for culture positive TB)[3], it represents an incremental performance improvement over previously described signatures. All of the blood-based signatures evaluated in this work (the ten-gene six-class signature, the threeGene signature and the ACS signature) show similar performance on the test datasets examined here, performance which falls below that observed with traditional sputum culture. While whole blood gene expression signatures do not appear likely to approach the performance of liquid culture, it is possible that whole blood signatures can be developed to improve diagnosis of TB cases who cannot produce sputum or who have paucibacillary disease, including HIV+ TB cases. Unfortunately, the lack of a “gold-standard” method of diagnosing TB when sputum culture cannot be obtained makes it extremely difficult to accurately evaluate blood transcriptional signatures in this context.

A possible practical application of this test is as a high-specificity “triage test” that can rule out patients unlikely to have TB, and identify persons who should receive a full sputum culture, thus reducing the necessity of working with difficult-to-acquire and potentially infectious sputum samples. At an operating point of 95% sensitivity, the ten-gene random forest shows a specificity of 47%. In a situation such as a medical clinic in a TB-endemic area, assuming 50% of patients presenting with symptoms consistent with TB have active TB, treating signature positive patients immediately would almost half the amount of sputum culture necessary.

## Supporting information

## List of Abbreviations

Mtb: Mycobacterium Tuberculosis;
AUC: Area under the receiver-operator curve;
ROC: receiver operator curve;
LOOCV: Leave-one-out cross-validation.
OD: other diseases;
TB: active tuberculosis;
LTB: latent tuberculosis;
ACS: Adolescent Cohort Study;
RF: random forest;
SVM: support vector machine;
NN: neural net;
NKN: k-nearest neighbours.

## Declarations

### Ethics approval and consent to participate

Not applicable

### Consent for publication

All authors have read and contributed to this manuscript, and approve of its publication

### Availability of data and material

All microarray datasets used in this study are available on GEO

### Competing interests

The authors declare that they have no competing interests

### Funding

This work was supported by the Strategic Health Innovation Partnerships (SHIP) Unit of the South African Medical Research Council, with funds received from the South African Department of Science and Technology. FJD was supported by the NCDIR, National Institutes of Health grant [U54 GM103511].

### Authors’ contributions

FJD, EGT, TJS and DEZ designed the analyses, assisted the interpretation of results, and revised the manuscript. FJD carried out the analysis and drafted the manuscript.

